# Viral outbreaks involve destabilized evolutionary networks: evidence from Ebola, Influenza and Zika

**DOI:** 10.1101/070102

**Authors:** Stéphane Aris-Brosou, Neke Ibeh, Jessica Noël

## Abstract

Recent history has provided us with one pandemic (Influenza A/H1N1) and two severe viral outbreaks (Ebola and Zika). In all three cases, post-hoc analyses have given us deep insights into what triggered these outbreaks, their timing, evolutionary dynamics, and phylogeography, but the genomic characteristics of outbreak viruses are still unclear. To address this outstanding question, we searched for a common denominator between these recent outbreaks, positing that the genome of outbreak viruses is in an unstable evolutionary state, while that of non-outbreak viruses is stabilized by a network of correlated substitutions. Here, we show that during regular epidemics, viral genomes are indeed stabilized by a dense network of weakly correlated sites, and that these networks disappear during pandemics and outbreaks when rates of evolution increase transiently. Post-pandemic, these evolutionary networks are progressively re-established. We finally show that destabilization is not caused by substitutions targeting epitopes, but more likely by changes in the environment *sensu lato*. Our results prompt for a new interpretation of pandemics as being associated with evolutionary destabilized viruses.

## Introduction

Over the past few years, humanity has been affected by three major zoonotic events, with an Influenza pandemic in 2009 [1], an Ebola virus outbreak in 2014-16 [2], and a Zika outbreak in 2015-16 [3]. In all these examples, the epidemiological and evolutionary dynamics of the pathogens involved, *i.e*., their phylodynamics [4], were meticulously reconstructed. For instance, in the case of Ebola, an initial phylogenetic study showed evidence that the outbreak originated from a single zoonotic event in an unknown animal reservoir [2], and that the resulting epidemic then spread to the largest and closest neighboring cities following the gravity model [5, 6], with some exceptions [7]. However, in this general context of severe outbreaks, we still do not quite fully understand what characterizes the evolutionary dynamics of the viruses during such events.

Recently, in an attempt to understand the genomic determinants of antigenic properties and drug resistance in influenza viruses, we described a novel algorithm to uncover pair of amino acids in a protein that evolve in a correlated manner [8]. We found that influenza A viruses show extensive evidence for correlated evolution, to such an extent that some amino acids evolve correlatively with more than one other site, hereby forming dense (undirected) networks (see also [9]). We furthermore uncovered that some of these pairs of sites are known to be epistatically interacting - specifically, experimental studies show that a mutation at one of these sites lowers viral fitness, which is then restored by a compensatory mutation [10]. Moreover, we showed that similar networks of sites can be found in the Ebola virus, with some of these sites also involved in episodes of adaptive evolution [11]. In light of these results, we here hypothesized that during an outbreak or a pandemic, these networks of tightly correlated sites might be transiently disrupted, hereby leading to a virus that is, from an evolutionary point of view, destabilized.

Such a network destabilization would require that some of the intrinsic properties describing these networks change in a similar manner across different viruses. One way of studying these properties is by resorting to the theory used in social networks analysis, and more generally developed in graph theory [12]. In our case, a network is made of nodes, that are amino acid sites in viral proteins, and a link between two amino acids means that these two sites show statistical evidence for evolving in a correlated manner. Both the structure of this network, and the pattern of connections among its nodes, influence its behavior: for instance, scale-free networks, where node connectivity follows a power law, are extremely robust to disruptions [13], just like dense networks [14], while the most connected nodes are also the most important ones in protein-protein interaction networks [15]. Such properties can be derived by summarizing a network with different statistics, such as the number of connections that a particular node has (its *degree*), or the shortest distance between each pair of nodes (the *average path length*).

In order to contrast the evolutionary dynamics of pandemic *versus* non-pandemic viruses, we here used these statistics to assess the stability of these networks of amino acids that evolve in a correlated manner. We predicted that viral evolutionary dynamics are weakened during a pandemic. As these dynamics often lead to complex networks of interactions [9, 11], we more specifically tested how the structure of these correlation networks is affected during an outbreak. We show that during a pandemic, the evolutionary dynamics of viral genes are severely disrupted, but also that they are progressively restored after the pandemic.

## Results

### Networks of correlated sites are destabilized during outbreaks

In search for evolutionary differences between regular epidemics and severe outbreaks, we first contrasted the glycoprotein precursor (GP) sequences of the Ebola virus that circulated before and during the 2014/2016 outbreak. For this, we identified with a Bayesian graphical model [16] the pairs of nucleotides that show evidence for correlated evolution in each data set, before and during the outbreak. As in previous work [9, 11], we found that these pairs of sites form a network. A first inspection of these networks of correlated sites revealed a striking difference between pre-2014 and outbreak sequences: in particular at weak correlations, the pre-2014 interaction networks are very dense and involve most sites of GP, while only a small number of sites are interacting in outbreak viruses (Figure 1). Furthermore, at increasing correlation strengths, outbreak networks become completely disconnected faster: at posterior probability *Pr* = 0.80 some sites still interact in pre-2014 proteins, while all interactions have disappeared from *Pr* = 0.60 in outbreak proteins (Figure 1). Similar patterns for the Influenza (at two antigenes, the hemagglutinin [HA] and the neuraminidase [NA]; Figures S3-S4) and Zika viruses (polymerase NS5; Figures S5) suggest that during a severe outbreak, an evolutionary destabilization of viral genes occurs, especially among sites that entertain weak interactions.

### Destabilization affects weakly correlated sites

To further investigate this destabilization hypothesis, we analyzed the structure of these networks with the tools of social network analysis and graph theory [12]. Again, we found a consistent pattern when contrasting regular and outbreak viruses: at weak to moderate interactions (*Pr* ≤ 0.50), outbreak viruses have networks of smaller diameter, shorter path length, and reduced eccentricity (Figure 2a-c, columns 1-5). All these patterns point to fewer connected sites in outbreak viruses. Betweenness is smaller for outbreak viruses (except Ebola), and transitivity tends to be larger (except Zika). These last two measures also suggest that interactions among sites are weakened in outbreak viruses. Other networks statistics failed to show a clear pattern (Figure S6): in particular, there were no clear differences in terms of degree, centrality or homophily - all properties that are not directly related to network stability.

### Post-outbreak re-stabilization

Should these weak interactions play a critical role in the stabilization of viruses outside of pandemics, we would expect to observe the strengthening of all network statistics as years go by after the pandemic. To test this prediction and estimate how long this re-stabilization process can take, we analyzed in a similar way all influenza seasons in the Northern hemisphere following the 2009 pandemic (until 2015-16). Consistent with our prediction, both HA and NA genes show a gradual transition between a typical pandemic state to a regular state in two-to-three seasons (Figure 2, column 5-6, respectively).

### Non-genetic sources of destabilization

To understand what the potential sources of this destabilization are, we assessed the involvement of viral antigenic determinants / epitopes. Should mutations accumulating in such epitopes be responsible for destabilization, we would expect (i) that weak interactions in non-pandemic viruses involve mostly epitopes, and (ii) that pandemics be associated with the disappearance of these interactions at epitopes first. Figure 3 shows no evidence supporting this hypothesis (*X*^2^ = 0.0663, *df* =1, *P* = 0.7967): non-pandemic viruses show a small number of predicted epitopes in their interaction network, that do not act as central hubs of these networks, while pandemic viruses may actually show an enrichment in interacting epitopes. This suggests that non-genetic factors are likely responsible for the initial destabilization of the genome of pandemic viruses. Changes in their ecology / environment (vector) cannot be ruled out.

## Discussion

To understand how evolutionary dynamics are affected during a viral outbreak, we compared non-outbreak and outbreak viruses. Based on the hypothesis that non-outbreak viruses are in a stable evolutionary equilibrium, and that such a stability is mediated by correlated evolution among pairs of sites in viral genes, we reconstructed the coevolution patterns in genes of non-outbreak and outbreak viruses. In line with our prediction, we found that outbreak viruses exhibit fewer coevolving sites than their non-outbreak counterparts, and that these interactions are gradually restored after the outbreak, at least in the case of the Influenza (2009 H1N1) virus for both HA and NA.

Two independent lines of evidence are consistent with our destabilization hypothesis. First, all three viruses showed temporary increases in their rate of molecular evolution during each outbreak [2, 3, 1]; such increases can be expected to disrupt the coevolutionary structure, and hence, destabilize viral genomes. We showed that epitopes were not particular targets of this mutational process. This can be expected, as mutations (i) most likely affect sites randomly, and (ii) are quickly lost from the viral population. Second, a probable cause of the epidemics can be identified in all cases studied here. For Influenza, the 2009 pandemic was caused by a series of reassortment events that affected the two genes studied here, HA (triple-reassortant swine) and NA (Eurasian avian-like swine) [1]. Such exchanges of segments can very well destabilize the evolutionary dynamics, at least of the implicated segments. Similarly, a zoonotic event was implicated in the Ebola outbreak [2], and a change of continent in the case of Zika [3, 17, 18]. These corresponding changes of environment *(sensu lato)* might have triggered the destabilizations observed here. In addition to such environmental changes, it is very likely that destabilization reflects a complex interaction between the genetics of viruses, their demographic fluctuations and environmental changes.

This argument is further supported by recent work in physics, where it was shown that dense networks are more resilient, *i.e*. resistant to small perturbations, than sparser ones [14]. Moreover, in their simplest example, these authors modeled abundances in a community of mutualistic species, where the mutualistic term describes the pairs of interacting species; perturbations were then applied to the system to assess resilience. They showed that small perturbations did not affect average abundances, which remained high - their ‘desirable’ state. However, above a particular perturbation threshold, a bifurcation occured and a new ‘undesirable’ state, at low abundances, was reached. Our results are consistent with a similar system behavior, where the network of correlated amino acids is resilient to perturbations up to a certain point, when a bifurcation to an ‘undesirable’ state (the pandemic) occurs, and the system returns to its resilient state. One major difference though is that we observed a progressive return to stability in the case of influenza, while the resilience model suggests a second bifurcation, *i.e*. an instantaneous change, to the ‘desirable’ state [14].

One outstanding question is about the importance of weak patterns of coevolution within a gene: how can it be explained that it is essentially weak correlations (around *Pr* = 0.25) that distinguish non-outbreak from outbreak viruses? In a recent study on mice, four phenotypes were quantitatively analyzed following large intercrosses, and linear regressions on pairs of quantitative trait loci were used to detect non-additive effects, *i.e*., epistasis; it was then shown that most epistatic interactions were weak and, critically, tended to stabilize phenotypes towards the mean of the population [19]. Viruses are not mice, and not all the correlations that we detected are involved in epistatic interactions, but both this work in mice and the evidence presented here go in the same direction: weak interactions have a stabilizing effect on viral genes and their phenotype (regular epidemics). It is further possible that the intricate nature of these weak correlation networks has higher-order effects [19], that in turn increase canalization and hence may help viruses weather modest environmental and genotypic fluctuations [20]. The elimination of these many weak interactions has a destabilizing effect that may be caused by or lead to outbreaks. Our findings call for a new interpretation of pandemics that, from an evo-lutionary point of view, appeared to be associated with unhealthy or diseased viruses. While the evidence shown here does not support the causal nature of this relationship, monitoring correlation networks could help forecast imminent outbreaks.

## Methods

### Sequence retrieval

Nucleotide sequences were retrieved for three viruses: Influenza A, Ebola, and Zika, for select protein-coding genes, chosen because they represent the most sequenced / studied genes for each of these viruses [11, 21, 22, 23]. All sequences were downloaded in May 2016 (Table S1).

Full-length Influenza A sequences were retrieved directly from the Influenza Virus Resource [24]. Only H1N1 sequences circulating in humans for the hemagglutinin (HA) and neuraminidase (NA) genes were downloaded. These two genes are also very commonly studied and largely sampled in public databases [22, 23]. Two types of data sets were constructed: one containing pandemic and non-pandemic sequences circulating in 2009, the pandemic year, and one containing pandemic sequences circulating from August 1 to July 31 of each season in the Northern temperate region between 2009/2010 and 2015/2016 (seven seasons in total). Only unique sequences were retrieved.

For Ebola, the virion spike glycoprotein precursor, GP, was retrieved because of its key role in the emergence of the 2014 outbreak showing evidence for both correlated and adaptive evolution [11] as follows. A GP sequence (KX121421) was drawn at random from the 2014 strain used previously [11] and was employed as a query for a BLASTn search [25] at the National Center for Biotechnology Information. A conservative *E*-value threshold of 0 (*E* < 10^−500^) was used, which led to 1,181 accession numbers. As most of these accession numbers correspond to full genomes, while only GP is of interest, we (i) retrieved all corresponding GenBank files, (ii) extracted coding sequences with ReadSeq [26] of all genes, (iii) concatenated the corresponding FASTA files into a single file, (iv) which was then used to format a sequence database for local BLASTn searches, and (v) used GP from KX121421 in a second round of BLASTn searches (*E* < 10^−250^, coverage > 75%).

In the case of Zika, sequences of 252 complete genomes were retrieved from the Virus Pathogen Resource (www.viprbrc.org). The RNA-dependent RNA polymerase NS5 was specifically extracted by performing local BLASTn searches as described above. It is one of the most studied Zika genes [21, 27], as it is essential for the replication of the virus [27].

### Phylogenetic analyses

Sequences were all aligned with Muscle [28] with the fastest options (-maxiters 1 -diags). Alignments were visually inspected with AliView [29] to remove rogue sequences and sequencing errors. Phylogenetic trees were inferred by maximum likelihood under the General Time-Reversible model with among-site rate variation [30] with FastTree [31]. As outbreak sequences (Ebola and Zika viruses) cluster away from non-pandemic sequences, we used the `subtreeplot()` function in APE [32] to retrieve accession numbers of pandemic sequences and hence separate them from non-pandemic sequences with minimal manual input. FastTree was used a second time to estimate phylogenetic trees of the subset alignments, with the same settings as above.

### Network analyses of correlated sites

Amino acid positions (“sites”) that evolve in a correlated manner were identified with the Bayesian graphical model (BGM) in SpiderMonkey [16] as implemented in HyPhy [33]. Briefly, ancestral mutational paths were first reconstructed under the MG94×HKY85 substitution model [34] along each branch of the tree estimated above at non-synonymous sites. These reconstructions were recoded as a binary matrix in which each row corresponds to a branch and each column to a site of the alignment. A BGM was then employed to identify which pairs of sites exhibit correlated patterns of substitutions. Each node of the BGM represents a site and the presence of an edge indicates the conditional dependence between two sites. Such dependence was estimated locally by a posterior probability. Based on the chain rule for Bayesian networks, such local posterior distributions were finally used to estimate the full joint posterior distribution [35]. A maximum of two parents per node was assumed to limit the complexity of the BGM. Posterior distributions were estimated with a Markov chain Monte Carlo sampler that was run for 10^5^ steps, with a burn-in period of 10,000 steps sampling every 1,000 steps for inference. Analyses were run in duplicate to test for convergence (Figures S1-S2).

The estimated BGM can be seen as a weighted network of coevolution among sites, where each posterior probability measures the strength of coevolution. Each probability threshold gives rise to a network whose topology can be analyzed based on a number of measures [12] borrowed from social network analysis and graph theory. We focused in particular on six statistics: average diameter, the length of the longest path between pairs of nodes; average betweenness, measures the importance of each node in their ability to connect to dense subnetworks; assortative degree, measures the extent to which nodes of similar degree are connected to each other (homophily); eccentricity, is the shortest path linking the most distant nodes in the network; average strength, rather than just count the number of connections of each node (degree), strength sums up the weights of all the adjacent nodes; average path length, measures the shortest distance between each pair of nodes. All measures were computed using the igraph R package ver. 1.0.1 [36]. Thresholds of posterior probabilities for correlated evolution ranged from 0.01 (weak) to 0.99 (strong). LOESS regressions were then fitted to the results.

### Epitope analyses

Epitopes were predicted using the NetCTL 1.2 Server [37]. Briefly, Cytotoxic T lymphocyte (CTL) epitopes are predicted based on a neural network algorithm trained on a database of human MHC class I ligands. Epitopes can be predicted for 12 MHC supertypes (A1, A2, A3, A24, A26, B7, B8, B27, B39, B44, B58, B62), that are broad families of very similar peptides for which independent neural network models have been generated. As such, we ran the epitope prediction for each supertype independently, on non-outbreak and outbreak viruses. Circos plots were generated with the circlize R package ver. 0.3.10 [38]. Scripts and sequence alignments used are available from github.com/sarisbro.

## Acknowledgements

We thank Jonathan Dench and George S. Long for discussions and comments, as well as two anonymous reviewers for additional comments. This work was supported by the Natural Sciences Research Council of Canada and by the Canada Foundation for Innovation (S.A.B.) and by the University of Ottawa (N.I., J.N.).

## Author information

Affiliations

**Department of Biology, University of Ottawa, Ottawa, ON K1N 6N5, Canada**

Stéphane Aris-Brosou, Neke Ibeh & Jessica Noël

**Department of Mathematics and Statistics, University of Ottawa, Ottawa, ON K1N 6N5, Canada**

Stéphane Aris-Brosou

## Contributions

S.A.B. designed the study, and wrote the paper. S.A.B., N.I. and J.N. performed research and analyses, and edited the paper. All authors approved the final version of the manuscript.

## Corresponding author

Stéphane Aris-Brosou.

## Supplementary information

1. Supplemental Information

**Figure 1. Correlation network of pre-outbreak and outbreak Ebola viruses**. Networks of correlated sites in the GP protein are shown in each panel. The top row shows networks for the viruses circulating before the 2014 outbreak (blue); the bottom row shows networks for outbreak viruses (red). Each column shows networks for different strengths of correlation, from weak (*Pr* = 0.05) to strong (*Pr* = 0.95). Nodes represent animo acid sites, and edges correlations. Node sizes are proportional to diameter.

**Figure 2. Network properties between pandemic and non-pandemic viruses**. Results are shown for Ebola (column 1), Zika (2) and Influenza viruses: for HA and NA circulating in 2009 in (3) and (4), respectively, and for pandemic viruses circulating between the 2009-10 (deep red) and the 2015-16 (deep blue) season in (5) and (6). Pandemic viruses are show in red, while non-pandemic ones are in blue. Shading: 95% confidence envelopes of the LOESS regressions. Five network measures are shown: (a) diameter, (b) average path length, (c) eccentricity, (d) betweenness, and (e) transitivity.

**Figure 3. Interacting residues in pandemic and non-pandemic viruses**. Results are shown for Ebola at weak correlations (*Pr* = 0.20). Coevolving positions in the alignment are identified with arabic numbers; for those that are predicted to be epitopes, supertypes (A1, A2, *etc*.) are shown.

